# An approximate derivate-based controller for regulating gene expression

**DOI:** 10.1101/579615

**Authors:** Saurabh Modi, Supravat Dey, Abhyudai Singh

**Author notes:** Departments of Electrical and Computer Engineering, Biomedical Engineering at the University of Delaware, Newark, DE 19716, USA.

## Abstract

Inside individual cells, protein population counts are subject to molecular noise due to low copy numbers and the inherent probabilistic nature of biochemical processes. Such random fluctuations in the level of a protein critically impact functioning of intracellular biological networks, and not surprisingly, cells encode diverse regulatory mechanisms to buffer noise. We investigate the effectiveness of proportional and derivative-based feedback controllers to suppress protein count fluctuations originating from two noise sources: bursty expression of the protein, and external disturbance in protein synthesis. Designs of biochemical reactions that function as proportional and derivative controllers are discussed, and the corresponding closed-loop system is analyzed for stochastic controller realizations. Our results show that proportional controllers are effective in buffering protein copy number fluctuations from both noise sources, but this noise suppression comes at the cost of reduced static sensitivity of the output to the input signal. Next, we discuss the design of a coupled feedforward-feedback biochemical circuit that approximately functions as a derivate controller. Analysis using both analytical methods and Monte Carlo simulations reveals that this derivative controller effectively buffers output fluctuations from bursty stochastic expression, while maintaining the static input-output sensitivity of the open-loop system. As expected, the derivative controller performs poorly in terms of rejecting external disturbances. In summary, this study provides a systematic stochastic analysis of biochemical controllers, and paves the way for their synthetic design and implementation to minimize deleterious fluctuations in gene product levels.

## I. INTRODUCTION

Advances in single-cell technologies over the last decade have revealed striking differences between individual cells of the same population. For example, the level of a given protein can vary considerably across cells within a population, in spite of the fact that cells are identical clones of each other and are exposed to the same environment [1]–[6]. Such intercellular stochastic differences in gene expression patterns have tremendous consequences for biology and medicine [7]–[12], including stochastic cell-fate assignment [13]–[17], microbial bet hedging [18], [19], bacterial and cancer drug-resistance [20], [21].

Stochastic variations in the level of protein primarily arise from two main sources:

- Low-copy number fluctuations in underlying biomolecular components (genes, mRNA, proteins). Moreover, this shot noise is amplified by the fact that transcription of genes is not a continuous process but happens in sporadic bursts [22]–[26].
- External disturbances in the protein synthesis rate due to fluctuations in expression-related machinery (RNA polymerases, Ribosomes, etc.) or intercellular differences in cell-cycle stage/cell size [27]–[30].

Given these noise sources, cells encode diverse regulatory mechanisms to suppress stochasticity in the level of a protein around a set point. Perhaps the simplest example of this is a negative feedback loop, where the protein directly or indirectly inhibits its own synthesis [31]–[43]. Such naturally-occurring feedbacks have been shown to be key motifs in gene regulatory networks [44]. Furthermore, design of in-vitro/in-silico synthetic feedback system based on linear PID or nonlinear controllers is an intense area of current research [45]–[53]. In this contribution, we investigate design of biochemical circuits that function as *approximate* proportional and derivative-based controllers, and systematically investigate their effectiveness in buffering protein noise levels.

In Section II, we introduce an open-loop model of stochastic gene expression where the protein is expressed in random bursts, and its expression is impacted by an upstream noisy input (Fig. 1). We provide exact analytical formulas for the protein mean and noise levels in open loop. Section II also introduces the mathematical tools to be used throughout the paper for the analysis of stochastic dynamical systems. In Section III and IV, we discuss designs of nonlinear biochemical circuits that function as approximate proportional and derivate controllers, respectively. Given the nonlinearities introduced by feedback loops, we use the linear noise approximation method [54], [55] to investigate their noise suppression properties, and validate the results by performing exact stochastic simulations of the feedback system.

**Fig. 1.**
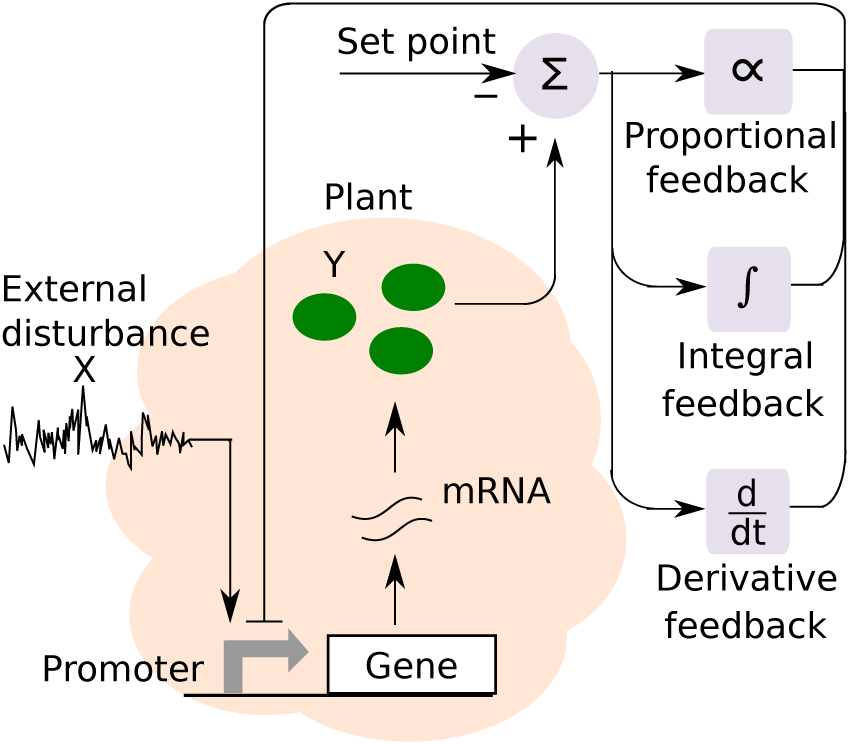
Schematic of the gene expression process, where a gene is transcribed to produce mRNAs. Each mRNA is subsequently translated to synthesize protein *Y* molecules. The expression process is impacted by an upstream external disturbance. Proportional-integral-derivative (PID) controllers can be designed to minimize fluctuations in *Y* copy number around a desired set point. This paper focuses on the design and stochastic analysis of proportional and derivative controllers, and integral feedback is omitted due to space constraints.

### Symbols and Notation

Throughout the paper we denote chemical species by capital letters, and use corresponding small letters for molecular counts. For example, if *Y* denotes a protein species, then *y*(*t*) is the number of molecules of *Y* at time *t* inside the cell. We use angular brackets to denote the expected value of random variables and stochastic processes. Given a scalar random process *y*(*t*) *∈* {0, 1, 2, *…*} that takes non-negative integer values, then

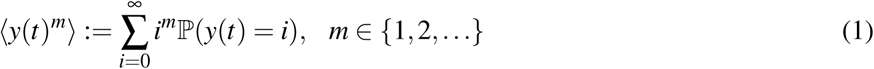

represent its *m*^*th*^ order uncentered moment and 𝕡(*y*(*t*) = *i*) is the probability of having *i* molecules. Steady-state statistical moments are denoted by

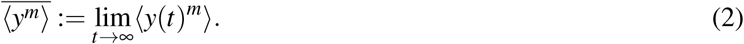

Finally, noise in the level of protein species is quantified by the steady-state coefficient of variation squared (variance divided by mean squared) that is defined as

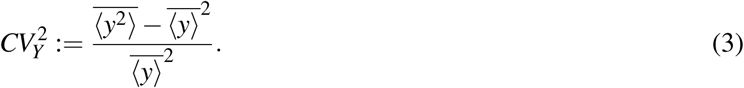

## II. SYSTEMS MODELING OF GENE EXPRESSION

We start by introducing simple models of the gene expression process with a particular focus on incorporating noise mechanisms that drive fluctuations in the level of a protein.

### A. Incorporating bursty dynamics

Transcription of individual genes inside single cells has been shown to occur in bursts of activity, followed by periods of silence [56]–[61]. Each burst corresponds to the gene stochastically switching to a transcriptionally active state, and then becoming inactive after synthesizing a few messenger RNA (mRNA) transcripts. These mRNAs are typically unstable with short half-lives, and each mRNA decays after translating a few protein molecules. The combined multiplicative effect of both these processes (single gene making multiple mRNAs, single mRNA making multiple proteins) is to create a net burst of protein molecules, every time the gene becomes active. Motivated by these experimental findings, we phenomenologically model protein copy-number fluctuations via a *bursty birth-death process* [62]–[68]. More specifically, bursts arrive at a constant Poisson rate *k*_*y*_ that corresponds to the frequency with which the gene becomes active. Each bursts arrival event, results in the synthesis of *B*_*y*_*∈*{1, 2, *…*} protein molecules, where the burst size *B*_*y*_ is an independent and identically distributed random variable that is drawn from an arbitrary positively-valued probability distribution.

Let *y*(*t*) denote the intracellular copy number of protein *Y* at time *t*. Then, based on the above model description, the probability of a burst event of size *B*_*y*_ = *j* molecules occurring in the next infinitesimal time interval (*t, t* + *dt*] is

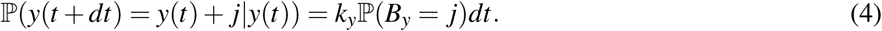

Assuming each protein molecule decays with a constant rate *γ*_*y*_, defines the probability for the protein death event occurring in the time interval (*t, t* + *dt*] as

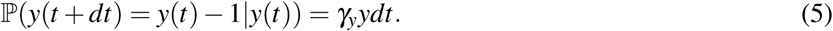

Having defined an integer-valued continuous-time Markov process *y*(*t*) via the probabilities (4)-(5), we now focus our attention on its statistical moments. We refer the reader to [69]–[72] for a thorough analysis of moment dynamics for stochastic systems of the form (4)-(5), and only provide the main result here – the time evolution of the expected value of *y*(*t*)^*m*^ is given by

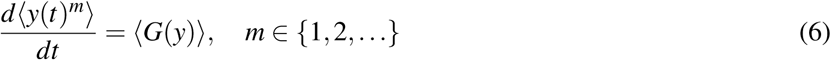

where the infinitesimal generator *G* takes the form

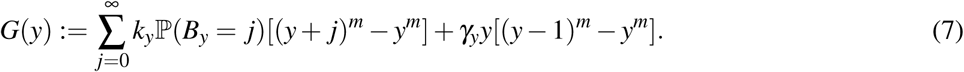

Intuitively, the right-hand-side of (7) is simply the product of the change in *y*^*m*^ when an event occurs and the probabilistic rate at which it occurs, summed across all possible events. Substituting the appropriate value of *m* in (6) yields the following moment dynamics

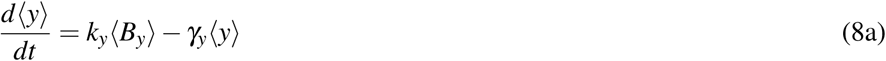

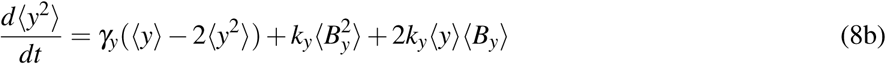

where ⟨*B*_*y*_⟩ is the mean protein burst size, and 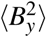 is its second-order moment. Subsequent steady-analysis of (8) reveals the protein mean and noise levels to be

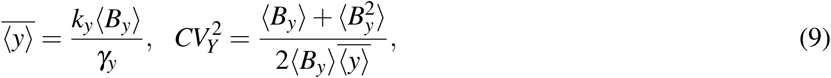

respectively. *B*_*y*_ = 1 with probability one leads to Poissonian fluctuations in *Y* copy numbers with 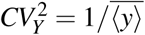. If the burst size *B*_*y*_ is assumed to be a geometrically-distributed random variable with mean burst size ⟨*B*_*y*_⟩ (as shown experimentally for an *E. coli* gene [73]), then 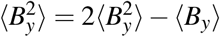, and the above noise levels reduce to

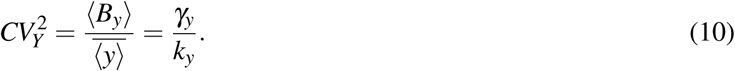

A key point worth mentioning is that the product 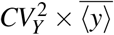 is independent of the burst frequency *k*_*y*_, while 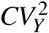 in (10) is independent of the mean burst size ⟨*B*_*y*_⟩. Thus, simultaneous measurements of both the mean and protein noise levels allows for discerning whether a change in 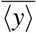 is a result of alterations in *k*_*y*_ or ⟨*B*_*y*_⟩. Interestingly, this noise-based method works quite well in practice, and has successfully elucidated the bursty kinetics of several genes [74]–[77].

### B. Incorporating external disturbance

Next, we introduce another important source of stochasticity that arises from external disturbances in the protein synthesis rate. These disturbances correspond to fluctuations in the abundance of enzymes, such as, transcription factors, RNA polymerases, etc. We lump these factors into a single species *X* and model its stochastic dynamics via a bursty birth-death process analogous to (4)-(5):

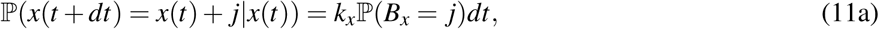

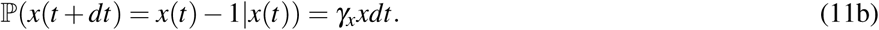

Here *k*_*x*_ is the arrival rate of bursts in *X*, *B*_*x*_ is the burst size, and *γ*_*x*_ is the decay rate of *X*. Then, as per (9)

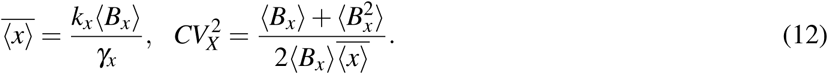

The disturbance is connected to the synthesis of *Y* by assuming that the frequency of protein *Y* bursts is proportional to *x*(*t*), and is given by 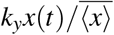. The division by 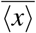 ensures that the average burst arrival rate is *k*_*y*_. This leads to a system of coupled bursty birth-death processes given by (11) and

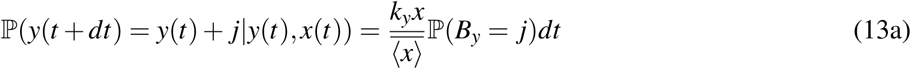

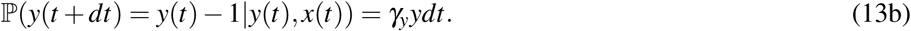

The statistical moments of this joint process evolve as per

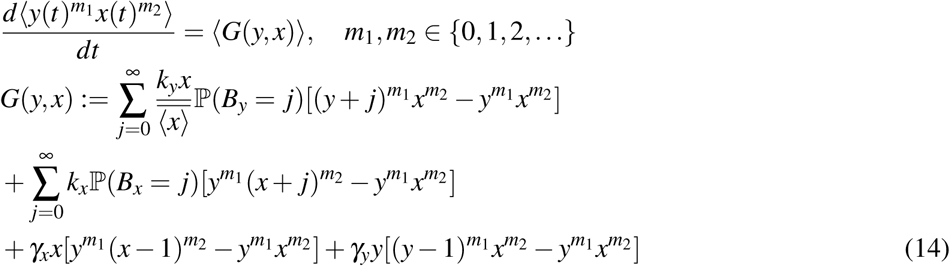

[69]–[72]. To write moment dynamics in a compact form we define a vector

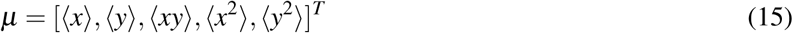

that consists of all the first and second order moments of *x*(*t*) and *y*(*t*). Then, its time evolution is given by a system of linear differential equations

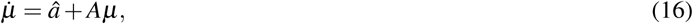

where vector *â* and matrix *A* are obtained via (15) by choosing appropriate values of *m*_1_, *m*_2_. Steady-state analysis of (16) results in the same mean *Y* level as (9), and the following noise level

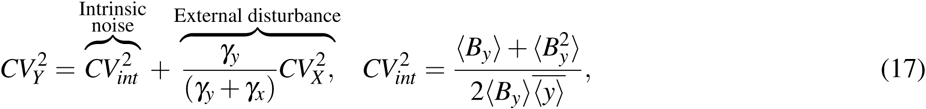

that can be decomposed into two components. The first component 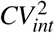 is the noise contribution from stochastic bursts computed earlier in (9), and has been referred to in literature as the *intrinsic noise* in *Y* [78]–[82]. The second component is the noise contribution of the external disturbance, and has been referred to as the *extrinsic noise* in *Y*. Note that the ratio *γ*_*y*_*/*(*γ*_*y*_ + *γ*_*x*_) quantifies the time-averaging of upstream fluctuation in *X* by *Y*. For example, fast fluctuations in *X* are efficiently averaged out by *Y*, and this ratio approaches zero for *γ*_*x*_*→*∞. In contrast, slow fluctuations in *X* lead to inefficient time-averaging that increases *Y* noise levels to

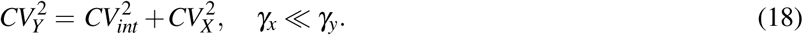

Next, we investigate how negative feedback regulation suppresses different noise components in (17) to minimize fluctuations in *Y* copy numbers around it mean 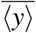.

## III. NOISE SUPPRESSION USING PROPORTIONAL CONTROLLER

To implement a negative feedback loop we first introduce a new protein species *Z* that functions as a noisy sensor of *Y*. Protein *Z* is also assumed to be synthesized in bursts of size *B*_*z*_, and senses *Y* via its burst frequency *k*_*z*_*y*(*t*) that responds linearly to any changes in *Y* levels. This leads to the following bursty birth-death process for *z*(*t*)

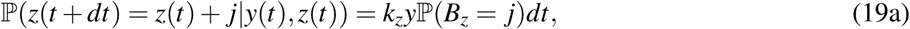

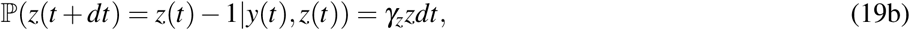

where *γ*_*z*_ is the decay rate of protein *Z*. Recall from Section II-B that the frequency of bursts in the *Y* protein was 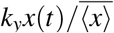 in the open-loop system. To close the feedback loop, we now modify this burst frequency to 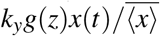, where *g*(*z*) is a positively-valued monotonically decreasing function of *z*(*t*). Typically, *g* takes the form of a Hill function that mechanistically arises from the fast binding-unbinding of the protein to the gene’s promoter region to regulate transcriptional activity [83]. Within this feedback there are three noise mechanisms at play: external disturbance *X* impacting synthesis of *Y*, expression of *Y* in stochastic bursts, and a noisy sensor *Z* that measures *Y* and inhibits it (Fig. 2). The overall stochastic system is given by (11), (19) and

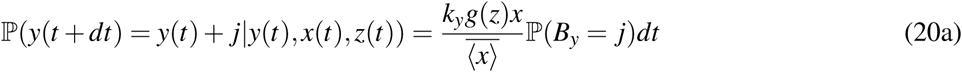

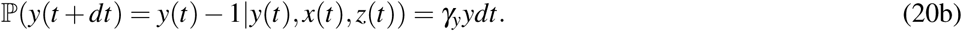

**Fig. 2.**
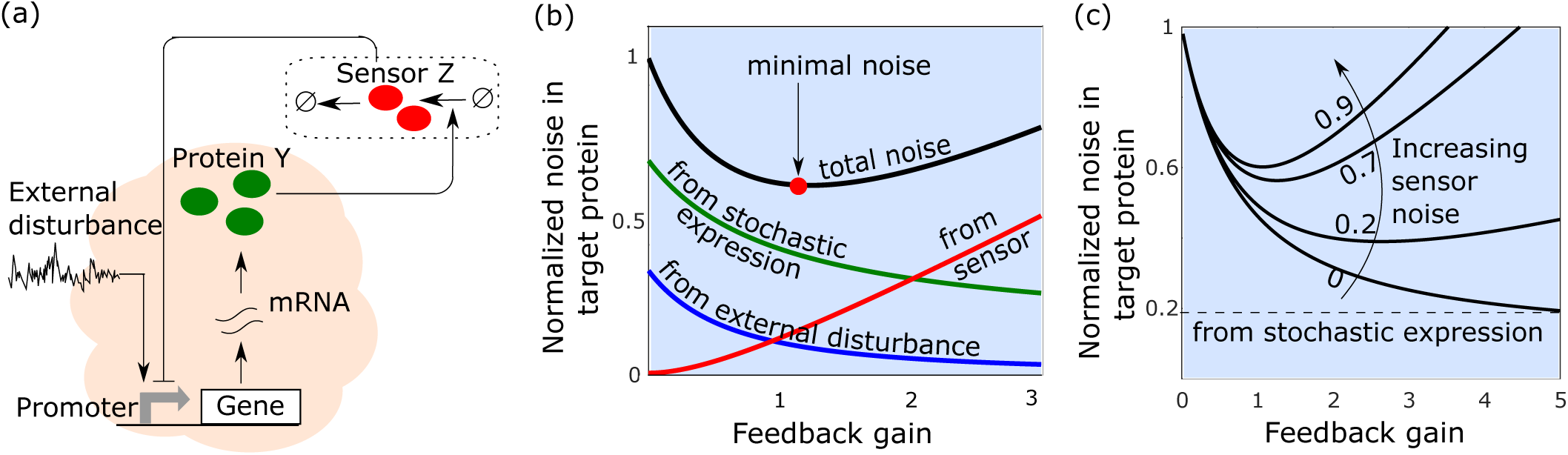
Implementation and noise decomposition for a proportional feedback controller. (a) Schematic of a proportional controller where the protein *Y* is sensed by a noisy sensor *Z* that inhibits the synthesis of *Y*. (b) Different components in the noise levels of protein *Y* from (28) plotted as a function of the feedback gain *f*_*p*_. While feedback selectively attenuates noise due to external disturbance and stochastic expression of *Y*, it amplifies the sensor noise, leading to a non-monotonic profile for the total noise. The noise contribution from the external disturbance decreases rapidly as a function of *f*_*p*_ and approaches zero for *f*_*p*_ *→*∞. In contrast, the intrinsic noise decreases slowly and asymptotically approaches a non-zero limit. In this plot, noise levels are normalized to the open-loop noise (*f*_*p*_ = 0), with other parameters chosen as 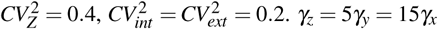. (c) The normalized total noise in *Y* from (28) with respect to the feedback gain *f*_*p*_ for different levels of sensor noise. The total noise 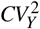 is minimized at an optimal feedback gain, which critically depends on the extent of sensor noise 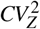.

### A. Analysis of Mean levels

At equilibrium, the mean levels of the random processes *x*(*t*), *y*(*t*) and *z*(*t*) satisfy

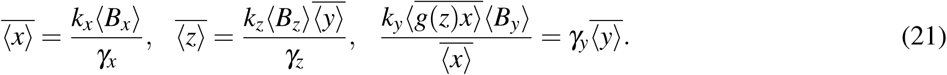

Assuming copy-number fluctuations are tightly regulated by the feedback system, and that they are small,

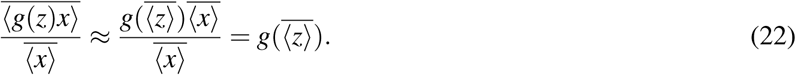

Given that *g*(*z*) is a positively-valued monotonically decreasing function, using (21) and (22), the steady-state mean level of *Y* is the unique solution to the equation

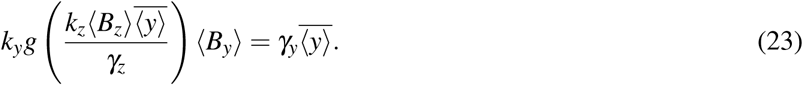

Having solved for the means, the burst frequency of *Y* can now be approximated using Taylor series as

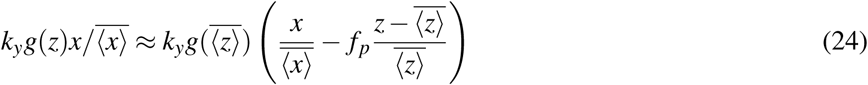

Where

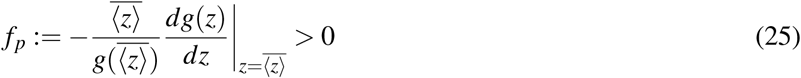

is the log sensitivity of the function *g* evaluated at steady state. Note that if the sensor dynamic is very fast compared to the measurand *Y* (i.e., *γ*_*z*_ *» γ*_*y*_), then *z*(*t*) ∝ *y*(*t*), and the burst frequency in (24) will be proportional to the error 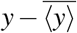. Hence, this circuit architecture can be interpreted as an approximate proportional controller with feedback gain *f*_*p*_. Finally, if we consider the parameter *k*_*y*_ in *Y* ‘s burst frequency as an environmental input, then one can define a static sensitivity of 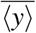 to *k*_*y*_

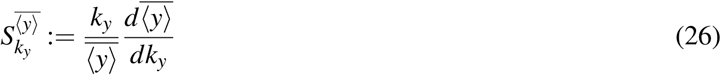

which using (21) and (25) is given by

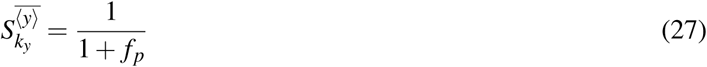

and monotonically decreases with increasing gain. Note for the open-loop system *f*_*p*_ = 0 and 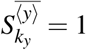 as mean 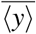 is simply proportional to *k*_*y*_ from (9).

### B. Analysis of Noise levels

Next, we focus on computing the noise levels in *Y* for the overall feedback system. As before, we define a vector *μ* that consists of all the first and second order moments of *x*(*t*), *y*(*t*) and *z*(*t*). The time evolution of *μ* can be obtained by expanding (15) to the three-species system, where *Y* ‘s nonlinear burst frequency is replaced by its linear approximation (24). Having linear probabilistic rates for all birth-death events results in a linear dynamical system (16) that can be solved analytically to obtain steady-state moments [72]. This analysis yields the following noise level for protein *Y*

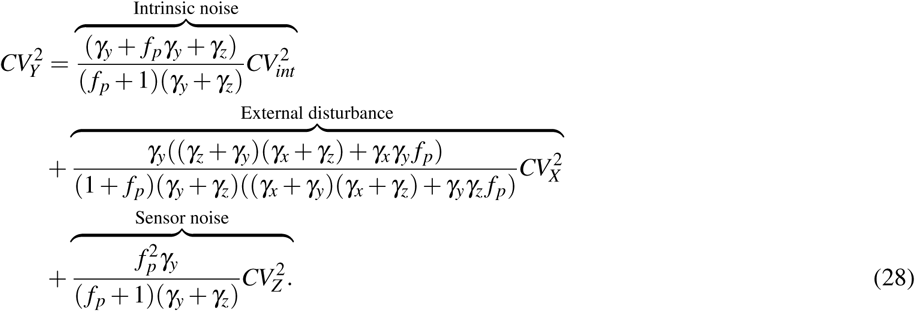

which can be decomposed into three components. The first component is the intrinsic noise in *Y* due to its bursty expression, and it decreases with increasing feedback gain *f*_*p*_ approaching a non-zero lower bound 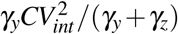 as *f*_*p*_ *→*∞. This lower bound represents a fundamental limit to which intrinsic noise can be decreased, and this limit is determined by how fast the sensor dynamics is compared to *Y* ‘s decay rate. The second component is the noise contribution from the external disturbance that monotonically decreases to zero as *f*_*p*_ *→*∞. The third component arises from the fact that the sensor *Z* is itself noisy, where

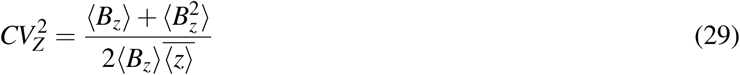

is the noise in *Z* due to its own expression occurring in random bursts. This third component is amplified with increasing feedback gain, and as a consequence, the total noise 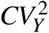 is a non-monotonic function of *f*_*p*_ with noise being minimal at an optimal feedback strength (Fig. 2). When *f*_*p*_ = 0, (28) reduces to the open-loop noise (17).

To further simplify the formula we assume that sensor dynamics is sufficiently fast (*γ*_*z*_ » *γ*_*y*_), and the time-scale of disturbance fluctuations are slow (*γ*_*x*_ « *γ*_*y*_). In this case, the sensor noise contribution becomes minimal, and (28) simplifies to

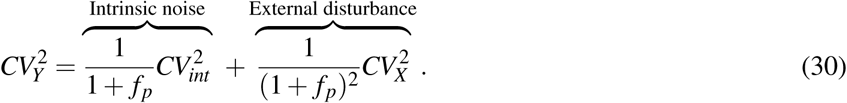

Note that the contribution from external disturbance decreases as 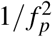 compared to 1*/ f*_*p*_ for the intrinsic noise. Hence, proportional feedback is much more effective in buffering stochasticity from external inputs rather than the intrinsic noise. This point relates to the static sensitivity 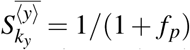 defined in (26), where increasing feedback gain suppresses noise, but it comes at the loss of adapting *Y* levels to changes in the environmental input.

## IV. NOISE SUPPRESSION USING DERIVATIVE CONTROLLER

Having completed the analysis for a proportional controller we next turn our attention to a derivative controller.

### A. Derivative controller design

How can biochemical circuits be designed to approximately sense the derivative of *y*(*t*)? To see this, consider the sensor dynamics in the deterministic limit

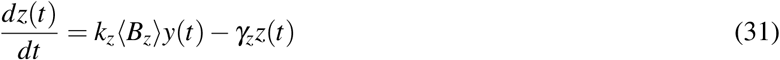

which in the Laplace domain corresponds to

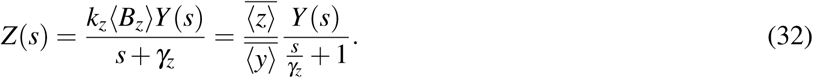

Here *Z*(*s*) and *Y* (*s*) are the Laplace transforms of *z*(*t*) and *y*(*t*), respectively, and we have used the fact that at equilibrium 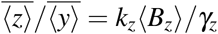. Now consider the scenario where *Z* activates the burst frequency of *Y*, and *Y* inhibits it own burst frequency (Fig. 3). This created an incoherent feedforward circuit that feedbacks into *Y*. Let the burst frequency of *Y* be proportional to (*z/y*)^*h*^, which corresponds to both *Y* and *Z* regulating the burst frequency with the same Hill coefficient *h*, and a strong binding affinity of the repressor *Y* [83], [84]. Then, in the limit of small fluctuations in *z*(*t*), *y*(*t*) around 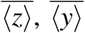, respectively,

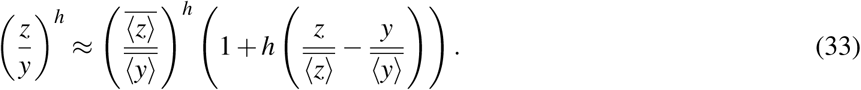

**Fig. 3.**
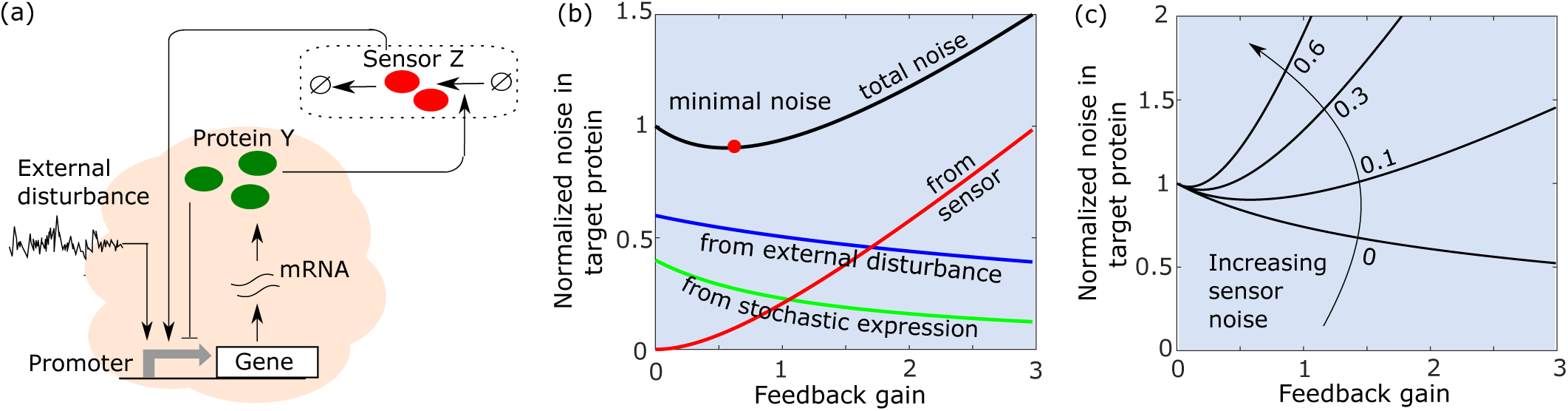
Implementation and noise decomposition for a derivative-based controller. (a) Schematic of the derivative controller where *Y* activates the sensor *Z, Z* activates the burst frequency of *Y*, while *Y* represses its own burst frequency (b) Different noise components in (39) are plotted as a function of the derivative feedback gain *f*_*d*_. Noise levels are normalized by the open-loop noise (17) and other parameters are chosen as 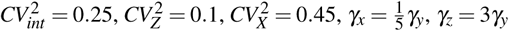.While both the intrinsic noise, and the noise contribution from the external disturbance decrease with increasing *f*_*d*_, the noise contribution from the sensor increases. In contrast to the proportional feedback, the intrinsic noise decreases faster than the disturbance contribution. (c) The noise in *Y* as a function of the derivative feedback gain *f*_*d*_ emphasizes the nonmonotonic noise profile for different levels of sensor noise 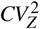.

Using (32) and assuming *γ*_*z*_ is large (i.e., fast sensor dynamics), the Laplace transform of the right-hand-side of (24) is

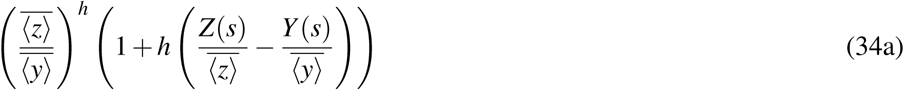

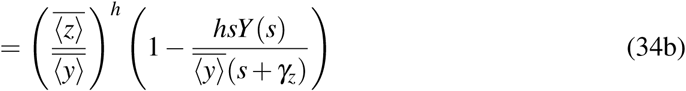

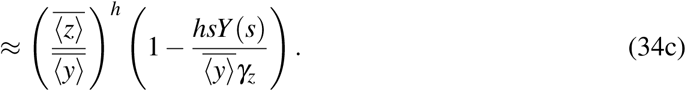

Recall that *sY* (*s*) is the Laplace transform of the time derivate of *y*(*t*), and hence in the time-domain, the burst frequency (24) corresponds to implementing a derivative controller. Going back to the original stochastic system, let the frequency of bursts in the *Y* protein be given by 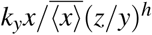. Then, as the ratio 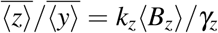 becomes a constant at steady-state, the mean protein level for *Y*

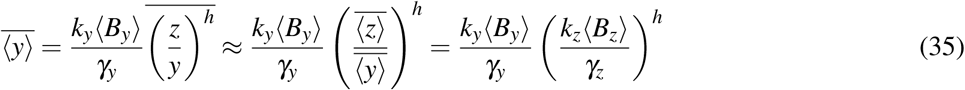

is proportional to *k*_*y*_ and the sensitivity 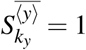 as in the open-loop system.

### B. Analysis of noise levels

To perform a noise analysis of the derivative controller-based feedback system, we revert to the noisy sensor *Z* described by the bursty birth-death process (19). The stochastic dynamics of protein *Y* is now described by

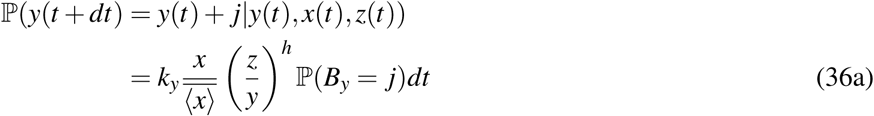

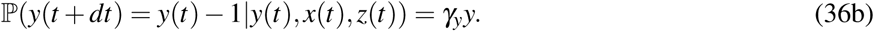

As before, the external disturbance is described by (11). To write a closed systems of differential equations for the time evolution of moments, we linearize protein *Y* ‘s burst frequency assuming small copy-number fluctuations

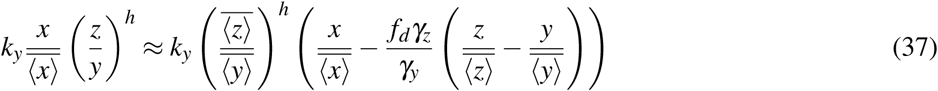

Where

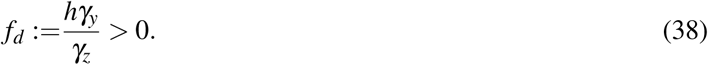

is the derivative feedback gain. A steady-state analysis of the resulting linear moment dynamics yields the following noise in protein *Y*

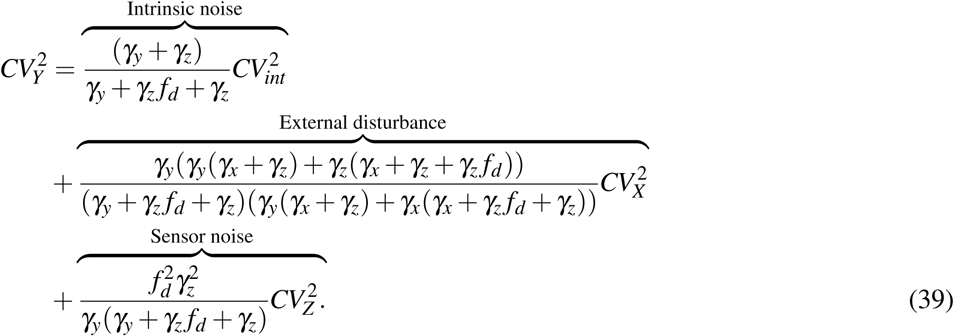

Analysis of the resulting noise components reveals that both the intrinsic noise, and the noise contribution from the external disturbance, decrease with increasing gain *f*_*d*_, with the former showing a much faster decay (Fig. 3). The noise contribution from the sensor amplifies with increasing feedback gain resulting in the total noise 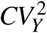 being minimized at an intermediate gain (Fig. 3). In the limit of slow fluctuations in the external disturbance *γ*_*x*_ « *γ*_*y*_, *γ*_*z*_, the above noise level simplifies to

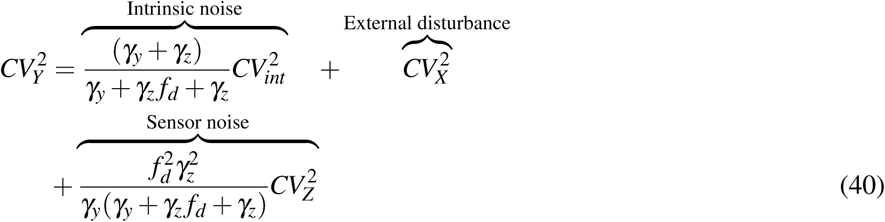

showing the derivative controller’s inability to reject low-frequency external disturbances. Finally, assuming no external disturbance 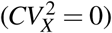, we verify the ability of a derivative controller to minimize intrinsic noise in *Y* by performing exact Monte Carlo simulations based on the Stochastic Simulation Algorithm (SSA) [85]. Stochastic simulation results of the overall nonlinear feedback system are shown in Fig. 4, and the noise levels show a good match with the formula (39) confirming the noise suppression abilities of a derivative controller.

**Fig. 4.**
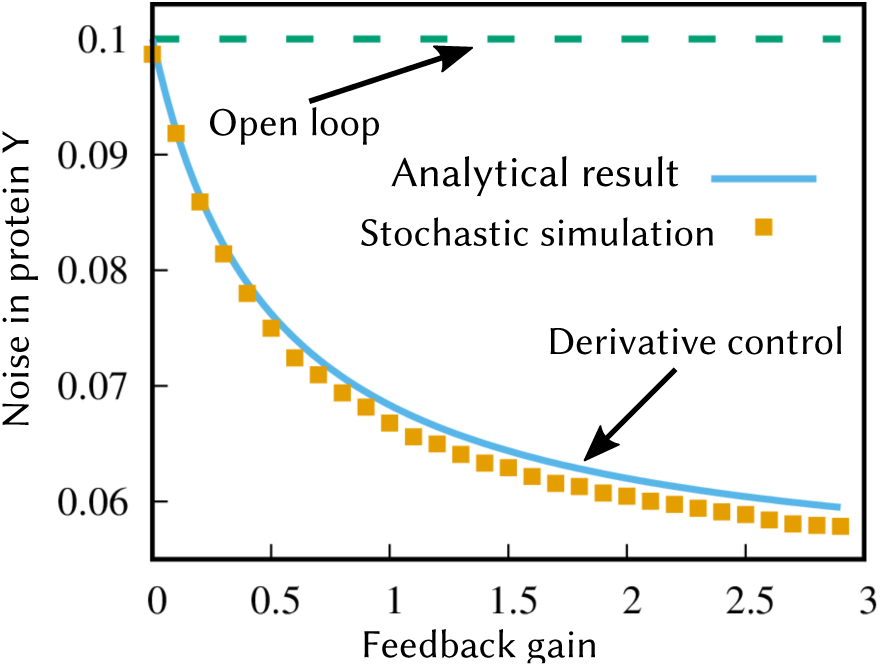
A derivative-based controller minimizes stochastic fluctuations in protein levels. Noise in the level of protein *Y* as obtained by performing exact Monte Carlo simulations of the nonlinear feedback system (19) and (36) with *h* = 1 and no external disturbance 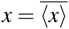 with probability one. Other parameters are taken as *B*_*y*_ = 20 with probability one, *k*_*y*_ = 2, *γ*_*y*_ = 0.2, *B*_*z*_ = 1 with probability one, *k*_*z*_ = *γ*_*z*_. In this case, *γ*_*z*_ was varied to change the gain *f*_*d*_ as per (38). The noise level obtained by running a large number of Monte Carlo simulations match their analytical estimates in (39).

## V. CONCLUSION

While PID controllers have become quite standard in industry, designing biochemical circuits that perform analogous functions inside cells is a highly nontrivial problem. Here we present simple circuits that function as *approximate* proportional and derivative controllers assuming fluctuations in molecular counts are small around their respective means. Our analysis of biochemically-implemented proportional feedback reveals the following properties:

- Proportional feedback is more efficient in suppressing stochasticity arising from noisy input signals, compared to noise arising from protein expression occurring in random bursts (Fig. 2).
- Any form of measurement noise (for example, due to stochastic expression of the sensor protein), leads to an optimal feedback gain for minimizing total protein noise, reminiscent of traditional feedback controllers.
- Noise suppression comes at the cost of reduced static input-output sensitivity, i.e., the protein levels are precisely regulated for a given environment, but do not respond to new environments.

We further provide design of a biochemical circuit for derivate-based control. In essence, the derivative of a signal is sensed by taking the difference of a delayed-signal (the sensor output) and the original signal. Intriguingly, our analysis shows that this controller suppresses intrinsic noise in the protein while preserving the open-loop static input-output sensitivity (Fig. 3). Intuitively, any rapid increase in protein levels due to a random burst is compensated by lowering the frequency of subsequent bursts. We confirmed our small-noise analysis results with exact Monte Carlo simulations of the nonlinear feedback system. As part of future work, we will investigate biochemical networks for integral feedback control, and constructing biological PID controllers for a given static input-output sensitivity, noise in the target protein, and transient response.

## ACKNOWLEDGMENT

AS is supported by the National Science Foundation Grant grant ECCS-1711548.

## REFERENCES

[1] J. M. Raser and E. K. O’Shea, “Noise in gene expression: Origins, consequences, and control,” Science, vol. 309, pp. 2010 – 2013, 2005.

[2] M. B. Elowitz, A. J. Levine, E. D. Siggia, and P. S. Swain, “Stochastic gene expression in a single cell,” Science, vol. 297, pp. 1183–1186, 2002.

[3] A. Bar-Even, J. Paulsson, N. Maheshri, M. Carmi, E. O’Shea, Y. Pilpel, and N. Barkai, “Noise in protein expression scales with natural protein abundance,” Nature Genetics, vol. 38, pp. 636–643, 2006.

[4] W. J. Blake, M. Kaern, C. R. Cantor, and J. J. Collins, “Noise in eukaryotic gene expression,” Nature, vol. 422, pp. 633–637, 2003.

[5] W. J. Blake, G. Balazsi, M. A. Kohanski, F. J. Isaacs, K. F. Murphy, Y. Kuang, C. R. Cantor, D. R. Walt, and J. J. Collins, “Phenotypic consequences of promoter-mediated transcriptional noise,” Molecular Cell, vol. 24, pp. 853–865, 2006.

[6] J. R. S. Newman, S. Ghaemmaghami, J. Ihmels, D. K. Breslow, M. Noble, J. L. DeRisi, and J. S. Weissman, “Single-cell proteomic analysis of S. cerevisiae reveals the architecture of biological noise,” Nature Genetics, vol. 441, pp. 840–846, 2006.

[7] E. Libby, T. J. Perkins, and P. S. Swain, “Noisy information processing through transcriptional regulation,” Proceedings of the National Academy of Sciences, vol. 104, pp. 7151–7156, 2007.

[8] D. L. Cook, A. N. Gerber, and S. J. Tapscott, “Modeling stochastic gene expression: implications for haploinsufficiency,” Proceedings of the National Academy of Sciences, vol. 95, pp. 15 641–15 646, 1998.

[9] R. Kemkemer, S. Schrank, W. Vogel, H. Gruler, and D. Kaufmann, “Increased noise as an effect of haploinsufficiency of the tumor-suppressor gene neurofibromatosis type 1 in vitro,” Proceedings of the National Academy of Sciences, vol. 99, pp. 13 783–13 788, 2002.

[10] A. Eldar and M. B. Elowitz, “Functional roles for noise in genetic circuits,” Nature, vol. 467, pp. 167–173, 2010.

[11] T. Wang and M. J. Dunlop, “Controlling and exploiting cell-to-cell variation in metabolic engineering,” Current Opinion in Biotechnology, vol. 57, pp. 10–16, 2019.

[12] A. H. K. Roeder, “Use it or average it: stochasticity in plant development,” Current Opinion in Plant Biology, vol. 41, pp. 8–15, 2018.

[13] A. Singh and L. S. Weinberger, “Stochastic gene expression as a molecular switch for viral latency,” Current Opinion in Microbiology, vol. 12, pp. 460–466, 2009.

[14] T. M. Norman, N. D. Lord, J. Paulsson, and R. Losick, “Memory and modularity in cell-fate decision making,” Nature, vol. 503, pp. 481–486.

[15] H. Maamar, A. Raj, and D. Dubnau, “Noise in gene expression determines cell fate in bacillus subtilis,” Science, vol. 317, pp. 526–529, 2007.

[16] G. Balázsi, A. van Oudenaarden, and J. J. Collins, “Cellular decision making and biological noise: From microbes to mammals,” Cell, vol. 144, pp. 910–925, 2014.

[17] H. H. Chang, M. Hemberg, M. Barahona, D. E. Ingber, and S. Huang, “Transcriptome-wide noise controls lineage choice in mammalian progenitor cells,” Nature, vol. 453, pp. 544–547, 2008.

[18] J.-W. Veening, W. K. Smits, and O. P. Kuipers, “Bistability, epigenetics, and bet-hedging in bacteria,” Annual Review of Microbiology, vol. 62, pp. 193–210, 2008.

[19] T. M. Norman, N. D. Lord, J. Paulsson, and R. Losick, “Stochastic switching of cell fate in microbes,” Annual Review of Microbiology, vol. 69, pp. 381–403, 2015.

[20] S. M. Shaffer, M. C. Dunagin, S. R. Torborg, E. A. Torre, B. Emert, C. Krepler, M. Beqiri, K. Sproesser, P. A. Brafford, M. Xiao, E. Eggan, I. N. Anastopoulos, C. A. Vargas-Garcia, A. Singh, K. L. Nathanson, M. Herlyn, and A. Raj, “Rare cell variability and drug-induced reprogramming as a mode of cancer drug resistance,” Nature, vol. 546, pp. 431–435, 2017.

[21] N. Balaban, J. Merrin, R. Chait, L. Kowalik, and S. Leibler, “Bacterial persistence as a phenotypic switch,” Science, vol. 305, pp. 1622–1625, 2004.

[22] D. Nicolas, B. Zoller, D. M. Suter, and F. Naef, “Modulation of transcriptional burst frequency by histone acetylation,” Proceedings of the National Academy of Sciences, vol. 115, pp. 7153–7158, 2018.

[23] O. Symmons, M. Chang, I. A. Mellis, J. M. Kalish, J. Park, K. Susztak, M. S. Bartolomei, and A. Raj, “Allele-specific rna imaging shows that allelic imbalances can arise in tissues through transcriptional bursting,” PLOS Genetics, vol. 15, p. e1007874, 2019.

[24] A. Raj, C. Peskin, D. Tranchina, D. Vargas, and S. Tyagi, “Stochastic mRNA synthesis in mammalian cells,” PLOS Biology, vol. 4, p. e309, 2006.

[25] S. Chong, C. Chen, H. Ge, and X. S. Xie, “Mechanism of Transcriptional Bursting in Bacteria,” Cell, vol. 158, pp. 314–326, 2014.

[26] D. M. Suter, N. Molina, D. Gatfield, K. Schneider, U. Schibler, and F. Naef, “Mammalian genes are transcribed with widely different bursting kinetics,” Science, vol. 332, pp. 472–474, 2011.

[27] O. Padovan-Merhar, G. P. Nair, A. G. Biaesch, A. Mayer, S. Scarfone, S. W. Foley, A. R. Wu, L. S. Churchman, A. Singh, and A. Raj, “Single mammalian cells compensate for differences in cellular volume and DNA copy number through independent global transcriptional mechanisms,” Molecular Cell, vol. 58, pp. 339–352, 2015.

[28] C. A. Vargas-Garcia, K. R. Ghusinga, and A. Singh, “Cell size control and gene expression homeostasis in single-cells,” Current opinion in systems biology, vol. 9, pp. 109–116, 2018.

[29] M. Soltani and A. Singh, “Effects of cell-cycle-dependent expression on random fluctuations in protein levels,” Royal Society Open Science, vol. 3, p. 160578, 2016.

[30] A. Mena, D. A. Medina, J. García-Martínez, V. Begley, A. Singh, S. Chávez, M. C. Muñoz-Centeno, and J. E. Pérez-Ortín, “Asymmetric cell division requires specific mechanisms for adjusting global transcription,” Nucleic Acids Research, vol. 45, pp. 12 401–12 412, 2017.

[31] A. Becskei and L. Serrano, “Engineering stability in gene networks by autoregulation,” Nature, vol. 405, pp. 590–593, 2000.

[32] Y. Dublanche, K. Michalodimitrakis, N. Kummerer, M. Foglierini, and L. Serrano, “Noise in transcription negative feedback loops: simulation and experimental analysis,” Molecular Systems Biology, vol. 2, p. 41, 2006.

[33] A. Singh and J. P. Hespanha, “Optimal feedback strength for noise suppression in autoregulatory gene networks,” Biophysical Journal, vol. 96, pp. 4013–4023, 2009.

[34] D. Orrell and H. Bolouri, “Control of internal and external noise in genetic regulatory networks,” Journal of Theoretical Biology, vol. 230, pp. 301–312, 2004.

[35] A. Borri, P. Palumbo, and A. Singh, “The impact of negative feedback in metabolic noise propagation,” IET Systems Biology, pp. 179–186, 2016.

[36] I. Lestas, G. Vinnicombegv, and J. Paulsson, “Fundamental limits on the suppression of molecular fluctuations,” Nature, vol. 467, pp. 174–178, 2010.

[37] Y. Tao, X. Zheng, and Y. Sun, “Effect of feedback regulation on stochastic gene expression,” Journal of Theoretical Biology, vol. 247, pp. 827–836, 2007.

[38] S. Kumar and A. J. Lopez, “Negative feedback regulation among sr splicing factors encoded by rbp1 and rbp1-like in drosophila,” The EMBO Journal, vol. 24, pp. 2646–2655, 2005.

[39] A. Singh and J. P. Hespanha, “Evolution of autoregulation in the presence of noise,” IET Systems Biology, vol. 3, pp. 368–378, 2009.

[40] D. J. Stekel and D. J. Jenkins, “Strong negative self regulation of prokaryotic transcription factors increases the intrinsic noise of protein expression,” BMC Systems Biology, vol. 2, p. 6, 2008.

[41] H. El-Samad and M. Khammash, “Regulated degradation is a mechanism for suppressing stochastic fluctuations in gene regulatory networks,” Biophysical Journal, vol. 90, pp. 3749–3761, 2006.

[42] P. S. Swain, “Efficient attenuation of stochasticity in gene expression through post-transcriptional control,” Journal of Molecular Biology, vol. 344, pp. 956–976, 2004.

[43] M. Voliotis and C. G. Bowsher, “The magnitude and colour of noise in genetic negative feedback systems,” Nucleic Acids Research, 2012.

[44] U. Alon, “Network motifs: theory and experimental approaches,” Nature Reviews Genetics, vol. 8, pp. 450–461, 2007.

[45] R. Sawlekar, F. Montefusco, V. V. Kulkarni, and D. G. Bates, “Implementing nonlinear feedback controllers using dna strand displacement reactions,” IEEE Transactions, vol. 15, p. 443, 2016.

[46] A. Milias-Argeitis, S. Summers, J. Stewart-Ornstein, I. Zuleta, D. Pincus, H. El-Samad, M. Khammash, and J. Lygeros, “In silico feedback for in vivo regulation of a gene expression circuit,” Nature Biotechnology, vol. 29, 2011.

[47] E. Klavins, “Proportional-integral control of stochastic gene regulatory networks,” IEEE conference on Decision and Control, 2010.

[48] G. Buzi and M. Khammash, “Implementation considerations, not topological differences, are the main determinants of noise suppression properties in feedback and incoherent feedforward circuits,” PLoS Computational Biology, vol. 12, p. e1004958, 2016.

[49] J. Uhlendorf, A. Miermont, T. Delaveau, G. Charvin, F. Fages, S. Bottani, G. Batt, and P. Hersen, “Long-term model predictive control of gene expression at the population and single-cell levels,” IET Systems Biology, vol. 109, 2012.

[50] C. Briat, C. Zechner, and M. Khammash, “Design of a synthetic integral feedback circuit: Dynamic analysis and dna implementation,” ACS Synthetic Biology, vol. 5, 2016.

[51] C. Briat, A. Gupta, and M. Khammash, “Antithetic integral feedback ensures robust perfect adaptation in noisy biomolecular networks,” Cell Systems, vol. 2, 2016.

[52] Y. Qian and D. D. Vecchio, “Realizing ‘integral control’ in living cells: how to overcome leaky integration due to dilution?” Journal of The Royal Society Interface, vol. 15, 2018.

[53] M. E. Wall, W. S. Hlavacek, and M. A. Savageau, “Design principles for regulator gene expression in a repressible gene circuit,” Journal of Molecular Biology, vol. 332, pp. 861–876, 2003.

[54] N. Van Kampen, Stochastic Processes in Physics and Chemistry. Elsevier. 2011.

[55] S. Modi, M. Soltani, and A. Singh, “Linear noise approximation for a class of piecewise deterministic markov processes,” American Control Conference (ACC), 2018.

[56] M. Chevalier, M. Gomez-Schiavon, A. Ng, and H. El-Samad, “Design and analysis of a proportional-integral-derivative controller with biological molecules,” bioRxiv, 2019, https://www.biorxiv.org/content/10.1101/303545v1.

[57] K. B. Halpern, S. Tanami, S. Landen, M. Chapal, L. Szlak, A. Hutzler, A. Nizhberg, and S. Itzkovitz, “Bursty gene expression in the intact mammalian liver,” Molecular Cell, vol. 58, pp. 147–156, 2015.

[58] A. M. Corrigan, E. Tunnacliffe, D. Cannon, and J. R. Chubb, “A continuum model of transcriptional bursting,” eLife, vol. 5, p. e13051, 2016.

[59] R. D. Dar, B. S. Razooky, A. Singh, T. V. Trimeloni, J. M. McCollum, C. D. Cox, M. L. Simpson, and L. S. Weinberger, “Transcriptional burst frequency and burst size are equally modulated across the human genome,” Proceedings of the National Academy of Sciences, vol. 109, pp. 17 454–17 459, 2012.

[60] A. Singh, B. S. Razooky, R. D. Dar, and L. S. Weinberger, “Dynamics of protein noise can distinguish between alternate sources of gene-expression variability,” Molecular Systems Biology, vol. 8, p. 607, 2012.

[61] T. Fukaya, B. Lim, and M. Levine, “Enhancer control of transcriptional bursting,” Cell, vol. 166, pp. 358–368, 2015.

[62] N. Kumar, A. Singh, and R. V. Kulkarni, “Transcriptional bursting in gene expression: Analytical results for genera stochastic models,” PLOS Computational Biology, vol. 11, p. e1004292, 2015.

[63] N. Kumar, T. Platini, and R. V. Kulkarni, “Exact Distributions for Stochastic Gene Expression Models with Bursting and Feedback,” Physical Review Letters, vol. 113, p. 268105, 2014.

[64] N. Friedman, L. Cai, and X. S. Xie, “Linking stochastic dynamics to population distribution: an analytical framework of gene expression,” Physical Review Letters, vol. 97, no. 16, p. 168302, 2006.

[65] V. Shahrezaei and P. S. Swain, “Analytical distributions for stochastic gene expression,” Proceedings of the National Academy of Sciences, vol. 105, pp. 17 256–17 261, 2008.

[66] K. R. Ghusinga, J. J. Dennehy, and A. Singh, “First-passage time approach to controlling noise in the timing of intracellular events,” Proceedings of the National Academy of Sciences, vol. 114, pp. 693–698, 2017.

[67] K. R. Ghusinga and A. Singh, “Effect of gene-expression bursts on stochastic timing of cellular events,” Proc. of the 2017 Amer. Control Conference, Seattle, WA, pp. 2118–2123, 2017.

[68] M. Soltani, C. Vargas, D. Antunes, and A. Singh, “Intercellular variability in protein levels from stochastic expression and noisy cell cycle processes,” PLoS Computational Biology, vol. 12(8), p. e1004972, 2016.

[69] J. P. Hespanha and A. Singh, “Stochastic models for chemically reacting systems using polynomial stochastic hybrid systems,” International Journal of Robust and Nonlinear Control, vol. 15, pp. 669–689, 2005.

[70] A. Singh and J. P. Hespanha, “Approximate moment dynamics for chemically reacting systems,” IEEE Transactions on Automatic Control, vol. 56, pp. 414–418, 2011.

[71] A. Singh, “Negative feedback through mRNA provides the best control of gene-expression noise,” IEEE transactions on nanobioscience, vol. 10, pp. 194–200, 2011.

[72] A. Singh and J. P. Hespanha, “Stochastic hybrid systems for studying biochemical processes,” Philosophical Transactions of the Royal Society A, vol. 368, pp. 4995–5011, 2010.

[73] J. Yu, J. Xiao, X. Ren, K. Lao, and X. S. Xie, “Probing gene expression in live cells, one protein molecule at a time,” Science, vol. 311, pp. 1600–1603, 2006.

[74] A. Singh, B. Razooky, C. D. Cox, M. L. Simpson, and L. S. Weinberger, “Transcriptional bursting from the HIV-1 promoter is a significant source of stochastic noise in HIV-1 gene expression,” Biophysical Journal, vol. 98, pp. L32–L34, 2010.

[75] E. M. Ozbudak, M. Thattai, I. Kurtser, A. D. Grossman, and A. van Oudenaarden, “Regulation of noise in the expression of a single gene,” Nature Genetics, vol. 31, pp. 69–73, 2002.

[76] R. D. Dar, S. M. Shaffer, A. Singh, B. S. Razooky, M. L. Simpson, A. Raj, and L. S. Weinberger, “Transcriptional bursting explains the noise–versus–mean relationship in mRNA and protein levels,” PLOS ONE, vol. 11, p. e0158298, 2016.

[77] R. Skupsky, J. C. Burnett, J. E. Foley, D. V. Schaffer, and A. P. Arkin, “HIV promoter integration site primarily modulates transcriptional burst size rather than frequency,” PLOS Computational Biology, vol. 6, p. e1000952, 2010.

[78] A. Singh and M. Soltani, “Quantifying intrinsic and extrinsic variability in stochastic gene expression models,” PLOS ONE, vol. 8, p. e84301, 2013.

[79] J. Paulsson, “Model of stochastic gene expression,” Physics of Life Reviews, vol. 2, pp. 157–175, 2005.

[80] A. Hilfinger and J. Paulsson, “Separating intrinsic from extrinsic fluctuations in dynamic biological systems,” Proceedings of the National Academy of Sciences, vol. 108, pp. 12 167–12 172, 2011.

[81] P. S. Swain, M. B. Elowitz, and E. D. Siggia, “Intrinsic and extrinsic contributions to stochasticity in gene expression,” Proceedings of the National Academy of Sciences, vol. 99, pp. 12 795–12 800, 2002.

[82] J. Paulsson, “Summing up the noise in gene networks,” vol. 427, pp. 415–418, 2004.

[83] U. Alon, An Introduction to Systems Biology: Design Principles of Biological Circuits. Chapman and Hall/CRC, 2011.

[84] N. Rosenfeld, M. B. Elowitz, and U. Alon., “Negative autoregulation speeds the response times of transcription networks.” Journal of Molecular Biology, vol. 323, pp. 785–793, 2002.

[85] D. T. Gillespie, “A general method for numerically simulating the stochastic time evolution of coupled chemical reactions,” Journal of Computational Physics, vol. 22, pp. 403–434, 1976.

